# *De novo* BK channel mutation causes epilepsy by regulating voltage-dependent, but not calcium-dependent, activation

**DOI:** 10.1101/120543

**Authors:** Xia Li, Sibylle Poschmann, Qiuyun Chen, Nelly Jouayed Oundjian, Francesca M. Snoeijen-Schouwenaars, Erik-Jan Kamsteeg, Marjolein Willemsen, Qing Kenneth Wang

## Abstract

Epilepsy is one of the most common neurological diseases. Here we report the first *de novo* mutation in the BK channel (p.N995S) that causes epilepsy in two independent families. The p.N995S mutant channel showed a markedly increased macroscopic potassium current mediated by increases in both channel open probability and channel open dwell time. Mutation p.N995S affects the voltage-activation pathway of BK channel, but does not affect the calcium sensitivity. Paxilline blocks potassium currents from both WT and mutant BK channels. We also identified two variants of unknown significance, p.E656A and p.N1159S in epilepsy patients. However, they do not affect BK channel functions, therefore, are unlikely to be a cause of disease. These results expand the BK channelopathy to a more common disease of epilepsy, suggest that the BK channel is a drug target for treatment of epilepsy, and highlight the importance of functional studies in the era of precision medicine.

## Introduction

Epilepsy is a group of neurological disorders and affects more than 65 million people worldwide (Thurman et al., 2011). Epilepsy is characterized by epileptic seizures and their neurobiological, cognitive, psychological, and social consequences. An epileptic seizure is a transient occurrence of signs or symptoms due to abnormally excessive or synchronous neuronal activities in the brain (Fisher et al., 2005). Epilepsy can be associated with other conditions such as stroke, brain trauma, and infection of the central of nervous system (acquired epilepsy) (Pandolfo, 2011). However, the causes of the majority of epilepsy are unknown (idiopathic epilepsy). Although extensive research has led to many interesting discoveries, the molecular mechanisms underlying the pathogenesis of epilepsy remain poorly understood.

One of the most exciting research areas in epilepsy has been the genetics in recent years. Genetic studies often uncover novel genetic and biological pathways for the pathogenesis of epilepsy. Further molecular characterization of the epilepsy genes identifies novel molecular mechanisms for the development of epilepsy. These studies further the understanding the pathophysiology of epilepsy, enable genetic testing and counseling, and may lead to development of new and optimal treatments. In 2005, our group identified a mutation in the *KCNMA1* gene that causes a human syndrome of coexistent generalized epilepsy and paroxysmal dyskinesia (GEPD) (Du et al., 2005). Seizures and epilepsy can be divided into two types dependent on the source of the seizures: either localized (focal seizure or epilepsy) or distributed (generalized seizure or epilepsy) within the brain (Chang and Lowenstein, 2003; Fisher, et al., 2005). The paroxysmal dyskinesia (PD) is a heterogeneous group of rare neurological conditions with recurrent episodes of sudden, unpredictable, disabling, and involuntary movement (Erro et al., 2014). *KCNMA1* encodes the large conductance voltage- and Ca^2+^-activated potassium channel α-subunit (also referred to as BK channel, calcium-activated potassium channel subunit α1, MaxiK channel, KCa1.1 or Slo1) (Horrigan and Aldrich, 2002).

The BK channel is expressed widely in various organs and tissues (McManus and Magleby, 1991), and plays an important role in numerous physiological processes such as repolarization of the membrane potential, control of neuronal excitability, neurotransmitter release, the control of smooth muscle tone, the tuning of hair cells in the cochlea and innate immunity (Brayden and Nelson, 1992; Fuchs and Murrow, 1992; Petersen and Maruyama, 1984; Robitaille and Charlton, 1992; Wu et al., 1999). The BK channel α-subunit contains seven transmembrane domains (S0-S6) at the N terminus with the voltage sensor S4 and an ionic pore with an activation gate between S5-S6 (Wallner et al., 1996), and a long C-terminal domain containing two regulator of conductance of K domains (RCK1 and RCK2) responsible for calcium sensing (Jiang et al., 2001; Quirk and Reinhart, 2001; Xia et al., 2002). Similar to many other voltage-activated potassium channels, the BK channel has a tetrameric structure, consisting of four pore-forming α-subunits. In addition to the pore-forming α-subunit, the diversity of the physiological properties of BK channels is generated by a series of auxiliary subunits comprising of β family and γ family of the BK channel accessory proteins (Behrens et al., 2000; Yan and Aldrich, 2012; Yan and Aldrich, 2010). The most interesting property of the BK channel is the dual synergistic activation pathway, including the voltage-dependent activation pathway and the calcium-dependent activation pathway, however the mechanisms of the channel gating are not fully known.

We previously studied a large family in which 16 family members developed epilepsy, paroxysmal nonkinesigenic dyskinesia (PNKD) or both and 13 unaffected family members (Du et al., 2005). Linkage analysis mapped the diseases gene on chromosome 10q22. Follow-up positional cloning identified a mutation in *KCNMA1* (c.1301G>A or p.D434G) which causes the disease in the family. The p.D434G mutation is located within the first RCK domain (RCK1) between transmembrane domains S6 and S8, which contains binding sites for regulatory ligands Ca^2+^ and Mg^2+^ (Shi et al., 2002; Xia et al., 2002). Our previous electrophysiological studies found that the p.D434G mutation in the BK channel leads to a gain of function. The mutation can increase the open probability of BK channels by enhancing the calcium sensitivity so that the BK channels with the p.D434G mutation can evoke a markedly greater macroscopic current than the wild-type (WT) BK channels under a physiological condition (Du et al., 2005; Yang et al., 2010).

In this study, we have identified a novel *de novo* mutation, p.N995S in the BK a-subunit, in the second RCK domain (RCK2) in two independent families with epilepsy alone. Electrophysiological studies revealed that the p.N995S mutation can increase macroscopic BK current density by increasing the open probability and the open duration of BK channels. However, the p.N995S mutation does not enhance the calcium sensitivity, thereby causing epilepsy by a novel mechanism. Our studies suggest that BK channel mutation can cause epilepsy alone and identify a novel molecular mechanism for epilepsy.

## Results

### Identification of an identical *de novo* mutation in *KCNMA1* in two independent families with epilepsy

Whole exome sequencing (WES) or targeted epilepsy panel sequencing with a panel of patients with epilepsy identified a heterozygous A to G transition in the *KCNMA1* gene (NIM Gene accession number z959149) in two independent patients with epilepsy (Figure 1A). The A to G transition results in the substitution of an asparagine residue for a serine residue at codon 995 (p.N995S) in the BK channel α-subunit. Both parents of the two patients are not affected (Figure 1A). The WES and targeted sequencing were performed for the patients and both parents, and the data indicate that both parents did not carry the p.N995S mutation (Figure 1A). Variant analysis from the sequencing validate the paternal relationship. Therefore, the p.N995S mutation occurred *de novo*.

**Figure 1.**
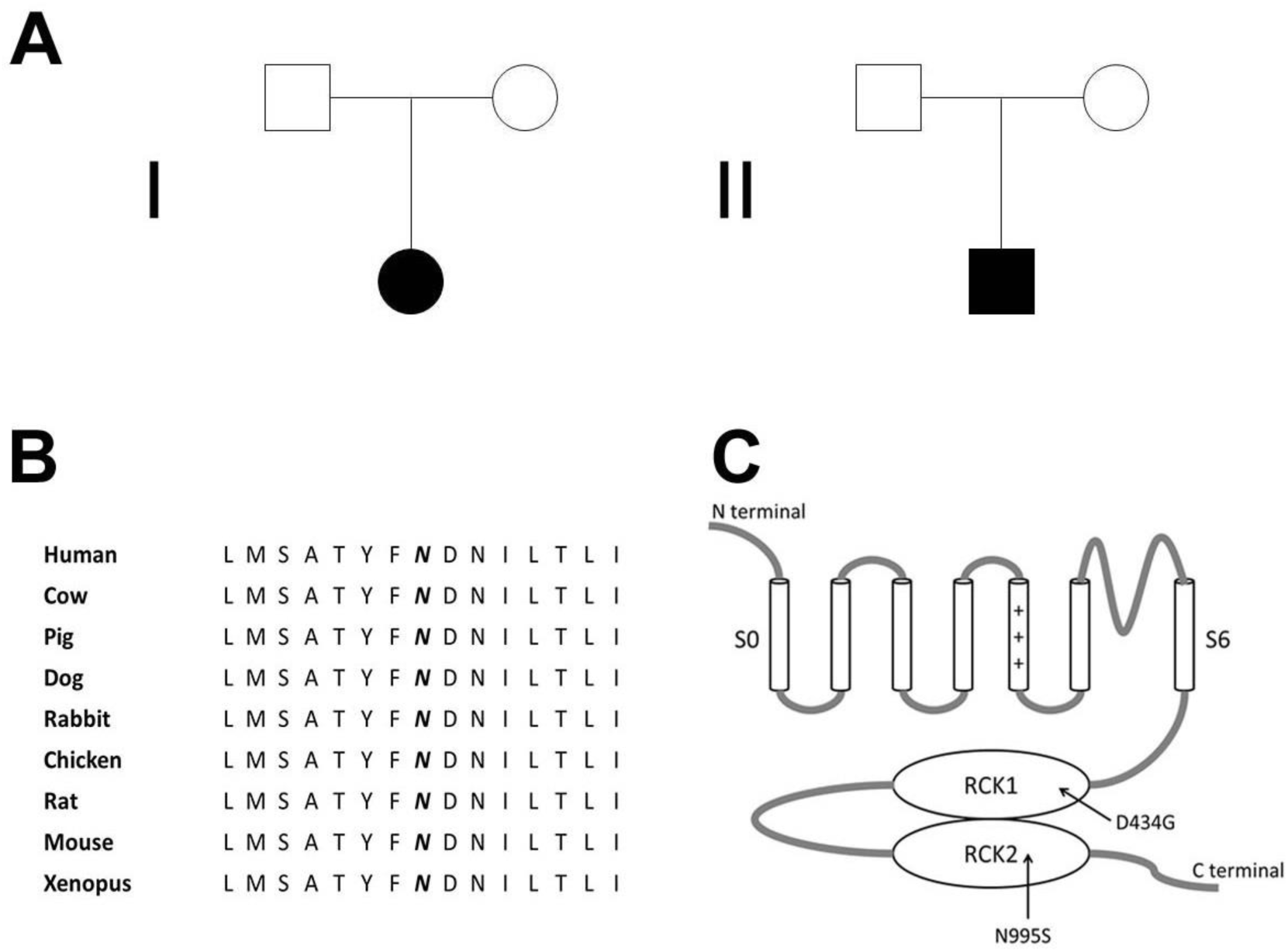
Whole exome sequencing and targeted epilepsy panel sequencing identified an identical *de novo* mutation p.N995S in the BK channel in two independent families. (A) Pedigrees of the two independent families. Squares represent males; circles, females; filled symbol, affected individual; open symbols, unaffected individuals. (B) The p.N995 residue, shown in bold and italic, of KCNMA1 is evolutionally conserved among different species. (C) Structure of the BK channel with the p.N995S mutation indicated. The p.D434G mutation identified previously in a large family with generalized epilepsy and paroxysmal dyskinesia is also shown.

The first epilepsy patient is a female from the Netherlands, who presented with neonatal convulsions. She is the oldest in a family with three younger healthy sisters born to consanguineous parents from Turkish ancestry. At the age of 9 years, her epilepsy was qualified as primary generalized myoclonic absence epilepsy. A developmental delay was present from the beginning. At the adult age, she had moderate to severe intellectual disability. She was able to speak in simple sentences. Under treatment with Levetiracetam, the epilepsy was under good control with only sporadic occurrence of seizures when tired. An EEG at the age of 17 years showed multifocal epileptic discharges. Brain MRI at the age of 17 years showed slight left sided insular white matter abnormalities. During physical examination at the age of 23 years, she had a short stature of 155 cm and small head circumference of 52.5 cm. She had no evident facial dysmorphic features. Neurological examination revealed no paroxysmal dyskinesia or other obvious abnormalities.

The second patient is also a female, but from Germany, who showed atypical absence epilepsy beginning at the age of 20 months. Semiology findings include arrest, mouth opening, head turning, variable altered consciousness, occasionally stronger seizures with risk of falls, duration of 10-20 sec, and frequency of > 20 per day. At the age of 3 years, she developed additional myoclonic seizures, which occurred 1-3 times per day. No paroxysmal dyskinesia was found. During febrile episodes the seizures improved. She was a couple of days seizure-free during varicella-zoster and streptococcus pyogenes infection. Her psychomotor development was mildly delayed. MRI scan and metabolic tests showed normal results. Her EEG showed generalized spike waves centrally and frontally located and photosensibility. Antiepileptic treatment with ethosuximide led to significant worsening of seizures, and oxcarbazepine and valproate showed also negative effects. Sultiame and perampanel did not influence the seizures. Levetiracetam reduced the seizure frequency and stopped myoclonic seizures. Zonisamide alone reduced the seizure frequency as well, but could not stop the myoclonic seizures. At the moment, she is treated with levetiracetam alone, her seizure activity is reduced at 10 per day, and the atypical absences are shorter. She currently retains consciousness, and shows no myoclonic seizures at the present time.

The p.N995S mutation occurs at a highly conserved amino acid residue among different species during evolution (Figure 1B). Moreover, the p.N995S mutation occurs within the second Ca^2+^-responsive RCK domain (Figure 1C). Identification of a *de novo* variant supports its pathogenicity. Identification of the same *do novo* mutation in two independent, unrelated families provides strong genetic evidence that the p.N995S mutation is highly likely to be pathogenic to epilepsy shared by the two families.

### The p.N995S mutation markedly increases the macroscopic potassium current from the BK channel

To determine whether the p.N995S mutation is a functional mutation that alters the function of the BK channel, we overexpressed the mutant channel and control wild type (WT) channel in HEK293 cells, respectively, and recorded their current-voltage relations. At an intracellular calcium concentration of 10 μΜ, the p.N995S mutant dramatically increased the activation rate (Figure 2B) and caused a large shift of the G-V curve to the negative potentials by 59 mV (Figure 2C). The data suggest that the mutant BK channel with the p.N995S mutation opens more easily and faster than the WT channel and that the p.N995S mutant channel generates a greater macroscopic BK current at a given calcium concentration and a membrane potential than the WT BK channel.

**Figure 2.**
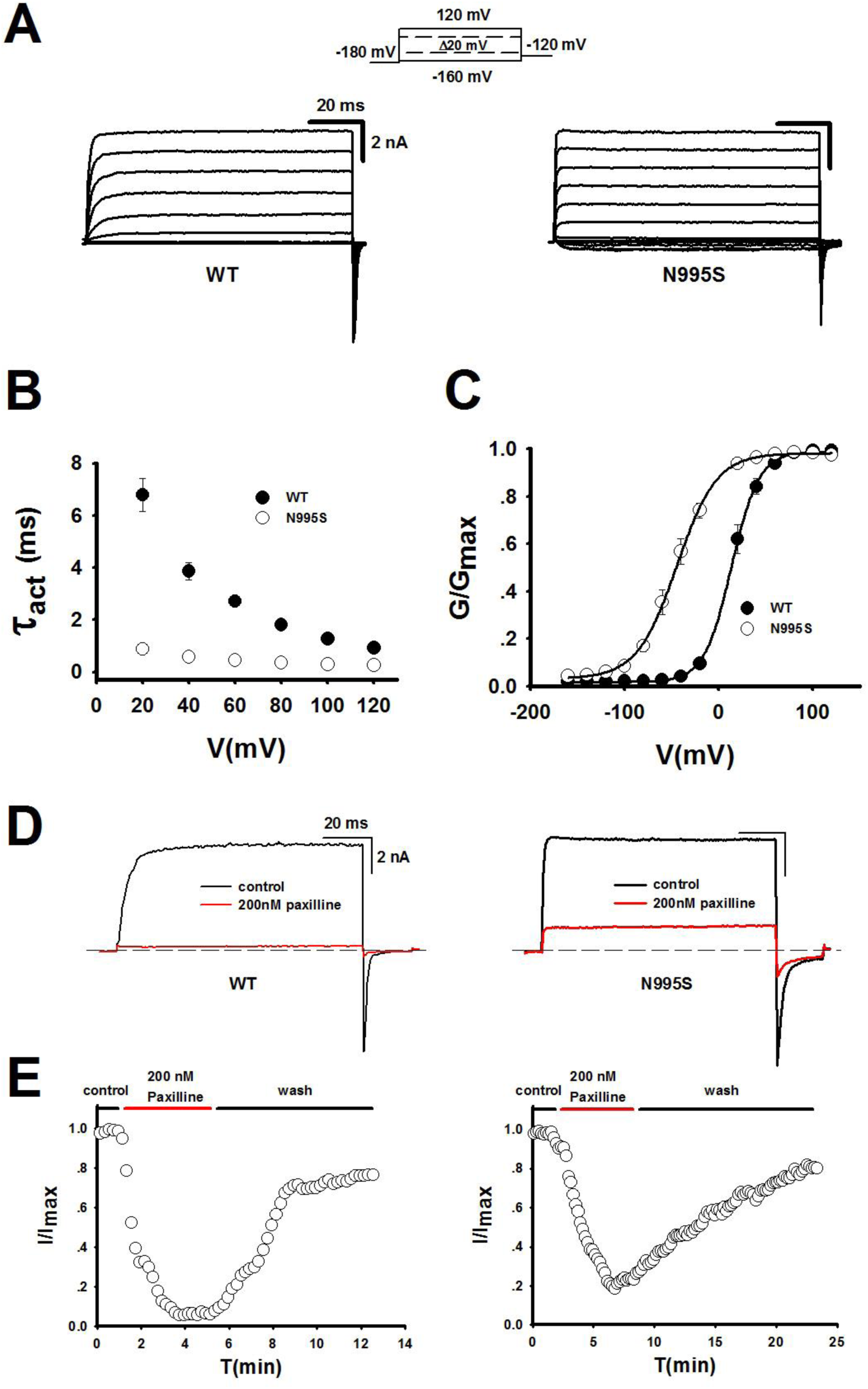
Electrophysiological characterization of WT and N995S mutant BK channels expressed in HEK293 cells. (A) Representative macroscopic currents of WT and N995S mutant BK channels from inside-out patches in the presence of 10 μΜ Ca^2+^ by the protocol as indicated. (B) Activation time constants of WT and N995S mutant BK channels at 10 μΜ Ca^2+^ are plotted against membrane potentials. (C) G-V curves of WT and N995S mutant BK channels at 10 μΜ Ca^2+^. All G-V curves are fitted by Boltzmann function (solid lines) with V1/2 (14.1 ± 6.6 mV for WT and -44.6 ± 8.2 mV for N995S). The data are presented as mean ± SD (n=6). (D) The representative traces of WT and N995S mutant BK channels from inside-out patches evoked by a test pulse 100 mV in the presence of 10 μΜ Ca^2+^ before (black) and after (red) application of 200 nM paxilline. (E) Normalized currents of WT (left) and N995S (right) mutant BK channels are plotted versus time. The black horizontal bar denotes the period without paxilline, whereas the red bar indicates the paxilline treat period.

Because the above data indicate that the p.N995S appears to be a gain-of-function mutation which significantly augments BK currents, BK channel inhibitors may block the potassium current and treat epilepsy. We examined the effect of paxilline, a selective BK-channel blocker, on the potassium currents generated by WT and p.N995S mutant BK channels. Treatment with 200 nM paxilline abolished the current from WT BK channel (Figure 2D and 2E). The p.N995S mutant BK channels are also sensitive to paxilline, but did not abolish the current (Figure 2D and 2E).

Because the β4 subunit of the BK channel is specifically expressed in the central nervous system (Brenner et al., 2005; Meera et al., 2000), we studied the effect of the epilepsy-linked p.N995S mutation on the BK channel in the presence of the β4 subunit. We characterized the current-voltage relations of the WT and mutant BK channels in the presence of the β4 subunit. The p.N995S mutation markedly enhanced activation of BK channels (Figure 3A) and shifted the G-V curve to a negative potential by 71 mV (Figure 3B) in the presence of the β4 subunit at a calcium concentration of 10 μΜ.

**Figure 3.**
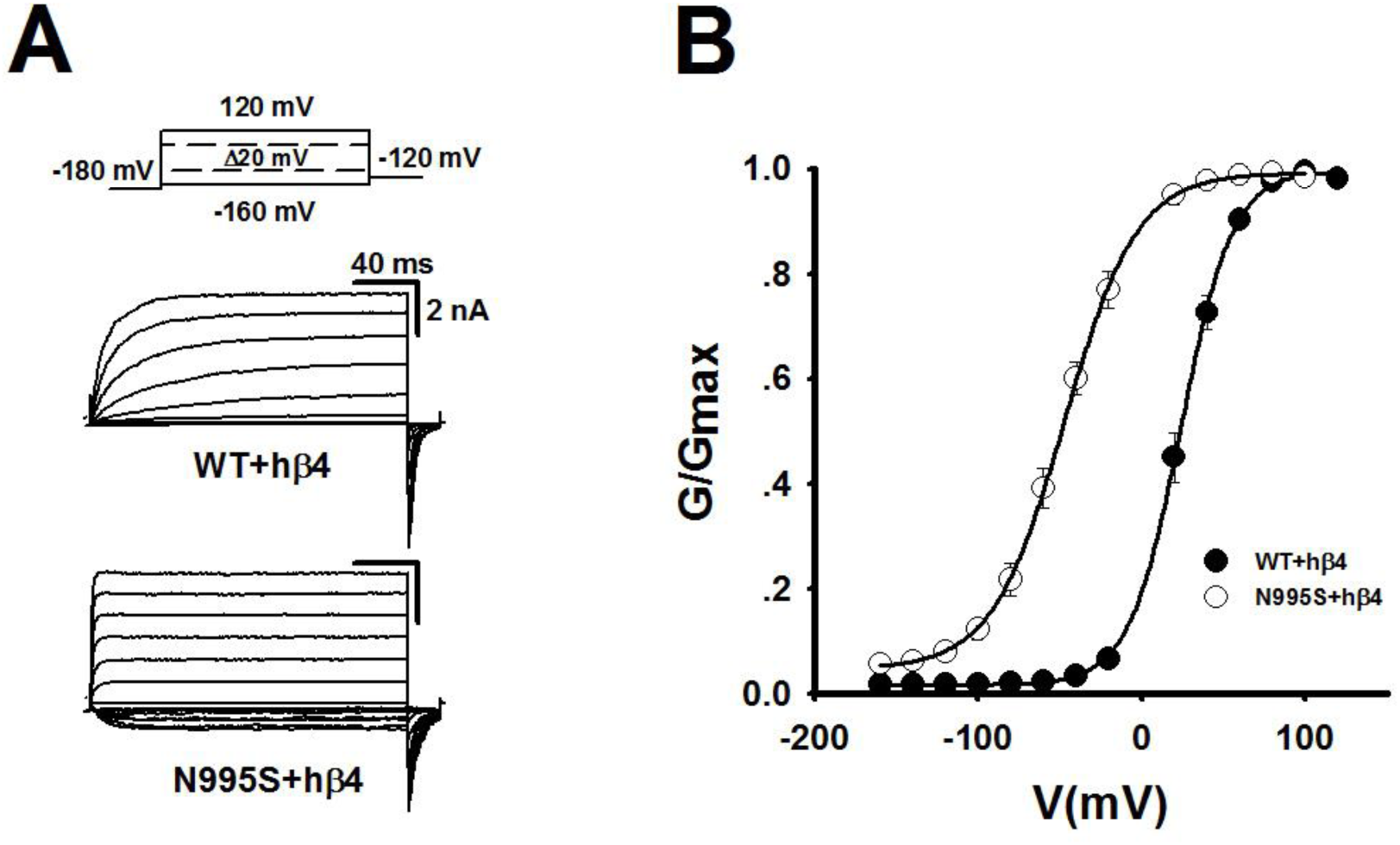
The β4 subunit slightly augments the effect of the N995S mutation on channel activation. (A) Representative macroscopic currents of WT and N995S mutant BK channels with hβ4 from inside-out patches in the presence of 10 μΜ Ca^2+^ by the protocol as indicated. (B) G-V curves of WT+hβ4 and N995S+hβ4 at 10 μΜ Ca^2+^. All G-V curves are fitted by Boltzmann function (solid lines) with V1/2 (24.1 ± 8.7 mV for WT+hβ4 and -47.1 ± 7.9 mV for N995S+hβ4). The data are presented as mean ± SD (n=6).

### The p.N995S mutation does not affect the calcium response sensitivity of the BK channel

The BK channel can be activated by both membrane potentials and intracellular calcium (Cui and Aldrich, 2000). Previously, we showed that the p.D434G mutation associated with GEPD increases the activation of the BK channel by enhancing calcium sensitivity (Du et al., 2005). Therefore, here we examined whether the p.N995S mutation associated with epilepsy alone increases the activation of the BK channel also by enhancing calcium sensitivity. We recorded the currents from WT and mutant BK channels under a series of calcium concentrations, including nominal 0 μΜ, 1 μΜ and 10 μΜ (Figure 4). The V_1/2_ values differed between the p.N995S mutant and WT channels at all the three different calcium concentrations (Figure 4C). Similarly, the absolute values of ΔV_1/2_ (V_1/2_ (N995S) – V_1/2_ (WT)) for the p.N995S mutant and WT channels were >40 mV at all the three different calcium concentrations (Figure 4B). However, the differences for the V_1/2_ values between the p.N995S mutant and WT channels remain similar at the three different calcium concentrations (Figure 4C). These results suggest that the p.N995S mutation increases BK channel activation not by enhancing calcium sensitivity. Moreover, the p.N995S mutant channel remained highly activated even at a nominal calcium concentration of 0 μΜ (Figure 4A and 4B).

**Figure 4.**
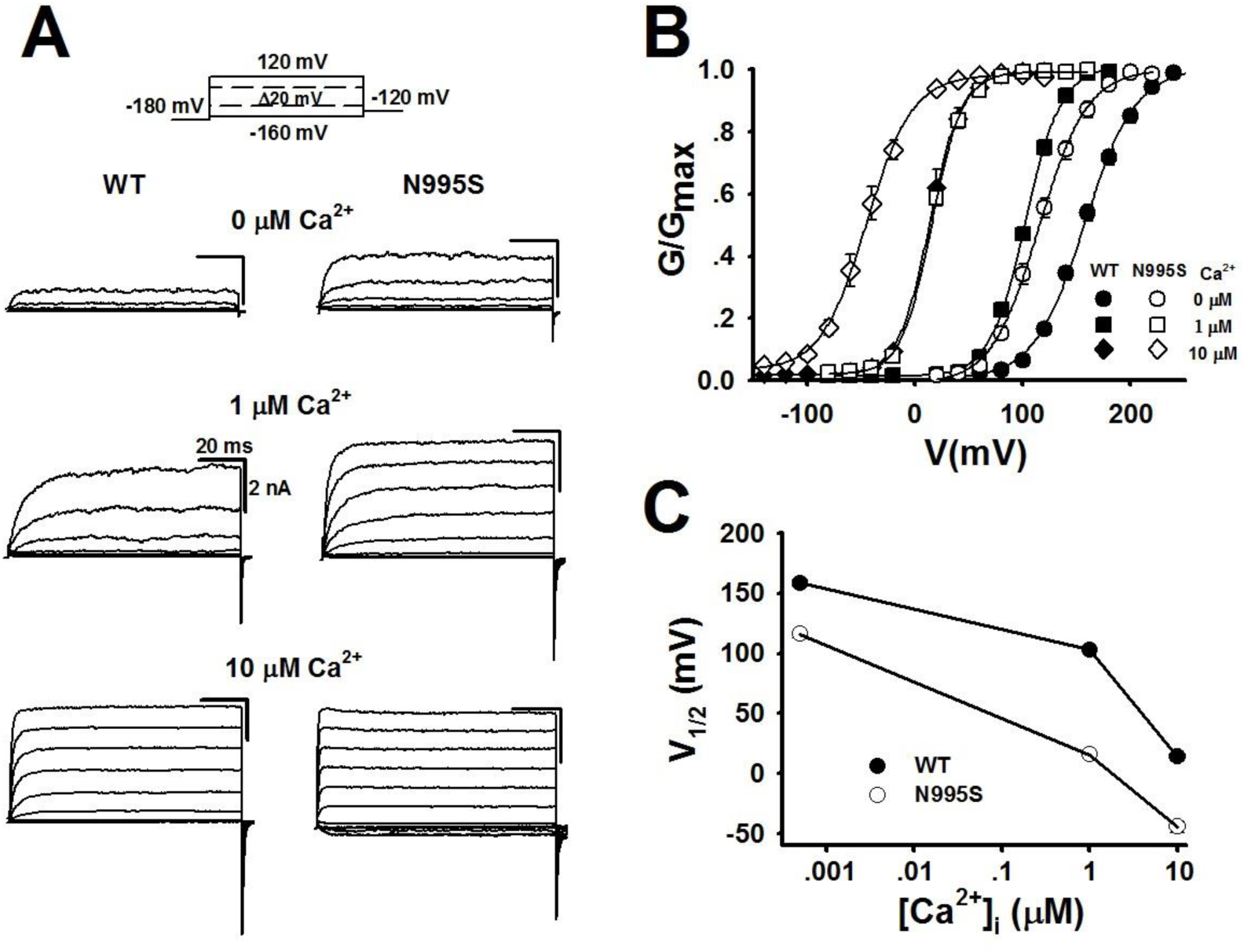
Effect of calcium concentrations on the function of N995S mutant BK channels. (A) Representative macroscopic currents of WT and N995S mutant BK channels from inside-out patches at nominal 0 μΜ Ca^2+^, 1 μΜ Ca^2+^ and 10 μΜ Ca^2+^ by the protocol as indicated. (B) G-V curves of WT and N995S mutant BK channels at nominal 0 μΜ Ca^2+^, 1 μΜ Ca^2+^ and 10 μΜ Ca^2+^. All G-V curves are fitted by Boltzmann function (solid lines) with V1/2 at nominal 0 μΜ Ca^2+^ (158.2 ± 6.3 mV for WT and 115.8 ± 6.9 mV for N995S), at 1 μΜ Ca^2+^ (102.8 ± 2.6 mV for WT and 15.8 ± 3.5 mV for N995S) and at 10 μΜ Ca^2+^ (14.1 ± 6.6 mV for WT and -44.6 ± 8.2 mV for N995S). (C) V1/2 of G-V curves versus [Ca^2+^]_i_ for WT and N995S mutant BK channels. The data are presented as mean ± SD (n=6).

### The p.N995S mutation continues to enhance activation of the channel even when the calcium response of the BK channel is blocked

As the calcium in the patch-clamp solutions cannot be eliminated completely, more evidence is needed to argue that the functional effect of the p.N995S mutation has little to do with the calcium activation pathway of BK channels. Therefore, we studied the effect of the p.N995S in mutant BK channels with mutations in the two putative calcium-binding sites, which are referred to as 2D2A (D427A, D432A) and 5D5N (D959N, D960N, D961N, D962N, D963N). Previous studies showed clearly that BK channels with 2D2A/5D5N mutations cannot be activated by calcium under low concentrations of calcium (Xia et al., 2002). As shown in Figure 5, the p.N995S mutant channels continued to generate greater macroscopic potassium currents than WT channels even when the two calcium-binding sites are mutated. Moreover, the G-V curves of the p.N995S mutant channel continued to shift to the negative potentials by more than 40 mV at nominal 0 μΜ, 1 μΜ, and 10 μΜ of calcium without a change of the slope as compared to the WT channel (Figure 5B). The differences of the V1/2 values between the p.N995S mutant channel and the WT channel were identical at nominal 0 μM, 1 μM, and 10 μM of calcium (Figure 5C) when the calcium-dependent activation pathway of BK channels is blocked by the 2D2A/5D5N mutations. Therefore, although the p.N995S mutation is located within the second RCK domain, its function is not related to the calcium sensitivity of the BK channel.

**Figure 5.**
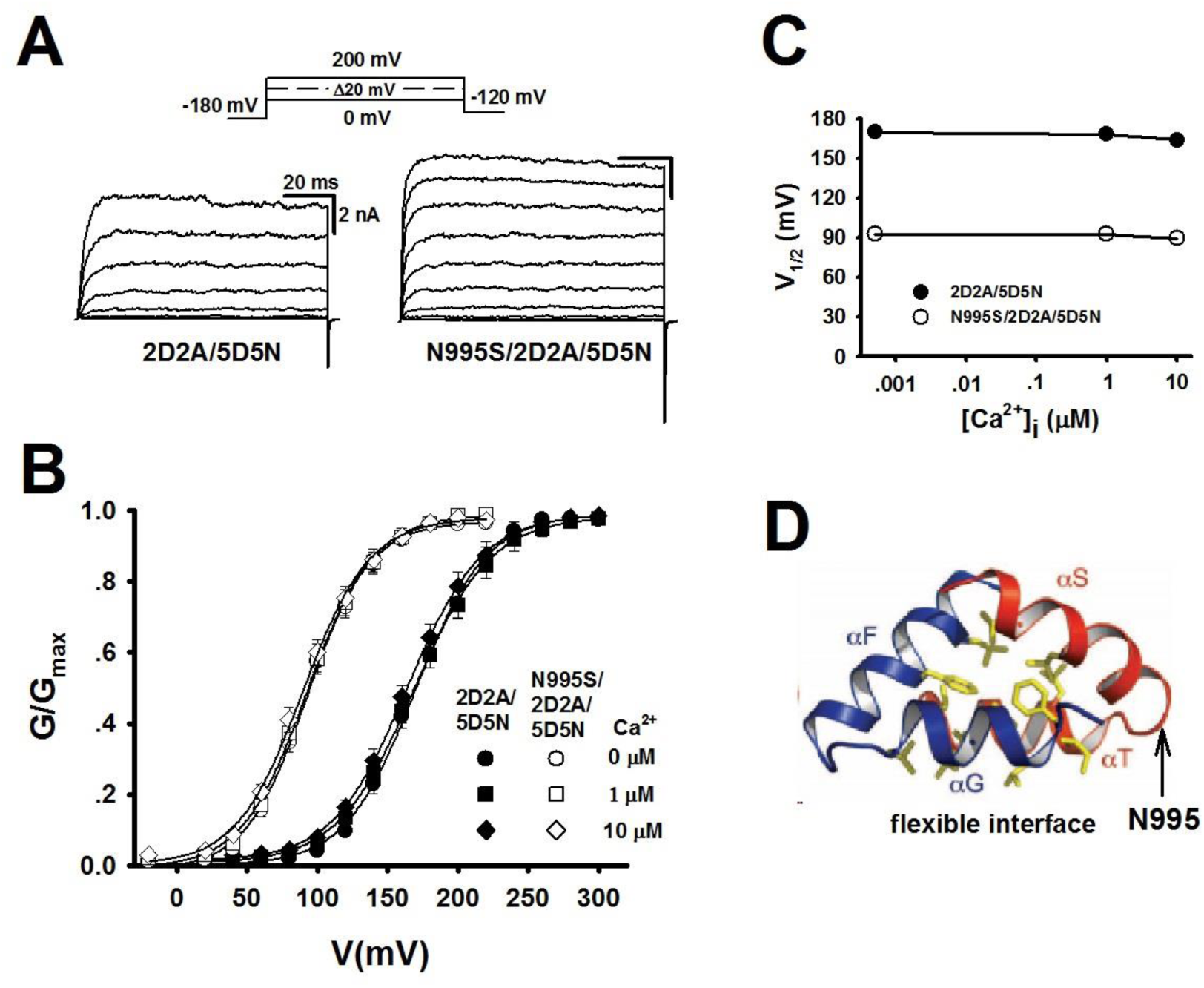
The functional effect of mutation N995S is not linked to the Ca^2+^-dependent activation mechanism of the BK channel. (A) Representative macroscopic currents of 2D2A/5D5N and N995S/2D2A/5D5N mutant BK channels from inside-out patches at nominal 0 μΜ Ca^2+^ by the protocol as indicated. (B) G-V curves of 2D2A/5D5N and N995S/2D2A/5D5N mutant BK channels at nominal 0 μΜ Ca^2+^, 1 μΜ Ca^2+^ and 10 μΜ Ca^2+^. All G-V curves are fitted by Boltzmann function (solid lines) with V1/2 and slope factor at nominal 0 μΜ Ca^2+^ (169.8 ± 7.1 mV, 24.6 ± 2.9 for 2D2A/5D5N and 92.8 ± 4.9 mV, 22.0 ± 2.6 for N995S/2D2A/5D5N), at 1 μΜ Ca^2+^ (168.6 ± 6.6 mV, 26.2 ± 3.0 for 2D2A/5D5N and 92.7 ± 5.1 mV, 23.2 ± 2.1 for N995S/2D2A/5D5N) and at 10 μΜ Ca^2+^ (164.0 ± 6.3 mV, 25.5 ± 3.2 for 2D2A/5D5N and 89.7 ± 5.0 mV, 23.6 ± 2.5 for N995S/2D2A/5D5N). (C) V1/2 of G-V curves versus [Ca^2+^]_i_ for 2D2A/5D5N and N995S/2D2A/5D5N mutant BK channels. (D) Schematic diagram showing that the N995S mutation is located within the flexible interface of the human BK channel [modified from Yuan et al] (Yuan, et al., 2010). The arrow indicates the location of the N995S mutation. The data are presented as mean ± SD (n=6).

### The p.N995S mutation stabilizes the open state of BK channels

To further investigate the functional impact of the p.N995S mutation on the BK channels, we analyzed its effects on single channel properties. Both the WT channel and the p.N995S mutant channel were activated by an increase in voltage at a given calcium concentration. The p.N995S mutant channels open more frequently than the WT channels (Figure 6A). The single channel conductance of the WT and mutant channels were 268.5±13.2 (pS), 276.3±18.9 (pS), respectively (Figure 6A). There is no statistically significant difference between the WT and mutant channels for the single channel conductance (*P*=0.260), which indicates that the p.N995S mutation does not affect the pore structure of the BK channel. The Po-V curve of the p.N995S mutant channels shifted to the negative potential at both 1 μM Ca^2+^ and 10 μM Ca^2+^ compared with the WT BK channels (Figure 6B), which is consistent with the data from the macroscopic current recordings (Figure 2). The distribution of the open dwell time was different between the WT channel and the mutant channel. The open dwell time histograms were fitted by an exponential function, and the time constant of the mutant channel was larger than that of the WT channel at the same potentials and calcium concentrations (Figure 6C and 6D). This suggests that the mutant channel stays open for a longer time in an open state compared with the WT channel. The open state of the mutant channel appears to be more stable than the WT channel. Thus, the single channel analysis identified the mechanism by which the p.N995S mutation enhances BK macroscopic currents, i.e. increased single-channel open probability and increased open dwell time.

**Figure 6.**
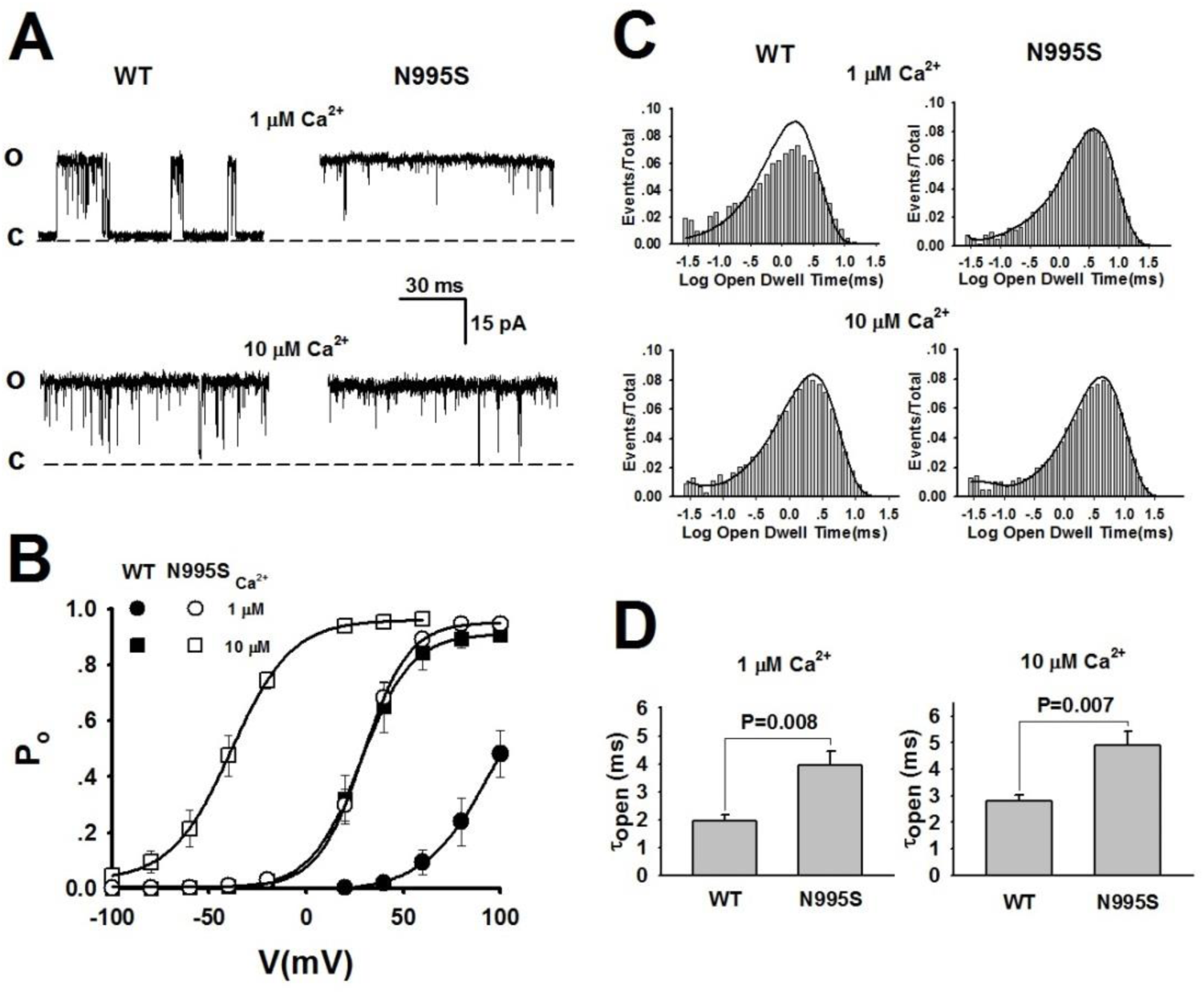
Single channel electrophysiological analysis of the N995S mutant BK channel. (A) Representative single channel currents of WT and N995S mutant BK channels at 1 μΜ Ca^2+^ and 10 μΜ Ca^2+^ recorded at 100 mV. C and O indicate the closed and open state of the channel, respectively. The sample rate is 50 kHz, and the low-pass filter is 5 kHz. (B) Open probability Po-V curves of WT and N995S mutant BK channels at 1 μΜ Ca^2+^ and 10 μΜ Ca^2+^. (C) Representative open dwell time histograms of WT and N995S mutant BK channels at 1 μΜ Ca^2+^ and 10 μΜ Ca^2+^ measured at 100 mV. Histograms are plotted in log-bin timescales and fitted by exponential function (solid lines). (D) Time constants of open dwell time for WT and N995S mutant BK channels at 1 μΜ Ca^2+^ (1.9 ± 0.2 ms for WT and 3.9 ± 0.5 ms for N995S) and at 10 μΜ Ca^2+^ (2.8 ± 0.2 ms for WT and 4.9 ± 0.5 ms for N995S). The data are presented as mean ± SEM (n=5).

### Identification of a benign variant p.N1159S in the BK channel in a patient with epilepsy and molecular characterization

Whole exome sequencing analysis identified a novel A to G transition at codon 1159 in a Chinese patient with epilepsy, which results in the substitution of the asparagine residue for a serine residue (p.N1159S). The p.N1159S occurs at an amino acid residue that is highly conserved during evolution (Figure 7B). The p.N1159S variant is located at the C-terminus of the BK channel (Figure 7C). This patient’s father also carried the heterozygous variant (Figure 7A). Both parents were not affected with epilepsy at the time of medical examination. However, low penetrance may occur and the father carrying the p.N1159S variant may not develop the disease at the time of diagnosis. Therefore, it is important to evaluate whether the p.N1159S variant is a functional mutation that increase predisposition to the disease or a benign variant.

We recorded the voltage-current relationship of the WT BK channel and the mutant channel with the p.N1159S variant (Figure 7D-G). No differences were observed between the mutant and WT channels with regard to the activation and current density of the BK channels in the presence or absence of the β4 subunit (Figure 7D-G). These data suggest that the p.N1159S variant is a benign variant that is unlikely to cause epilepsy in the family.

**Figure 7.**
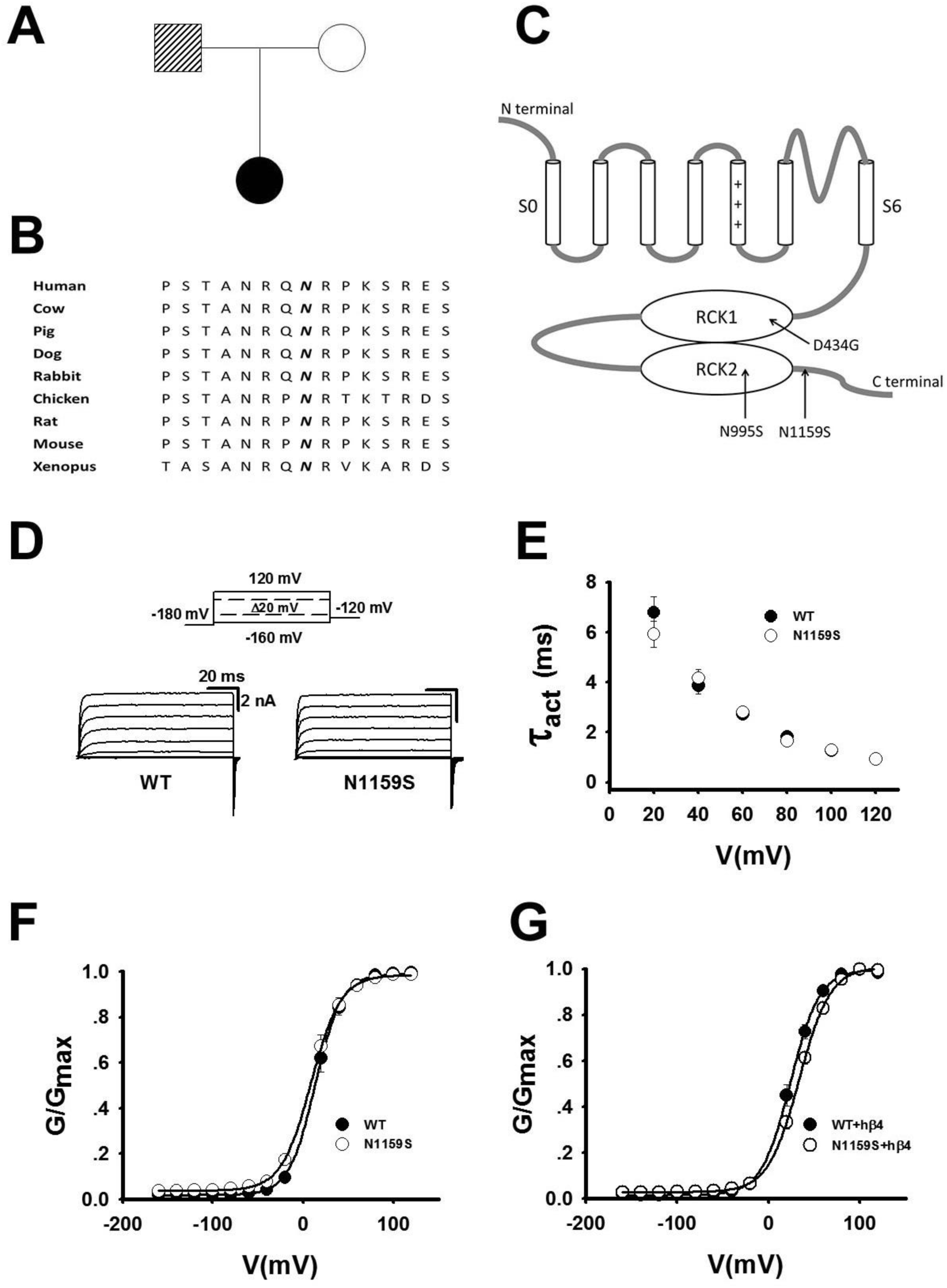
Whole exome sequencing identified a genomic variant N1159S in the BK channel in a family with epilepsy. (A) Pedigree of the family affected with epilepsy. Square, male family member; circles, females; filled symbol, affected individual; open symbols, unaffected individuals. (B) The N1159 residue, shown in bold and italic, of KCNMA1 is evolutionally conserved among different species. (C) Structure of the BK channel with the N1159S mutation indicated. D434G and N995S mutations are also shown. (D) Representative macroscopic currents of WT and N1159S mutant BK channels from inside-out patches in the presence of 10 μΜ Ca^2+^ by the protocol as indicated. (E) Activation time constants of WT and N1159S mutant BK channels at 10 μΜ Ca^2+^ are plotted against membrane potentials. (F) G-V curves of WT and N1159S mutant BK channels at 10 μΜ Ca^2+^. All G-V curves are fitted by Boltzmann function (solid lines) with V1/2 (14.1 ± 6.6 mV for WT and 9.3 ± 6.7 mV for N1159S). (G) G-V curves of WT+hβ4 and N1159S+hβ4 at 10 μΜ Ca^2+^. All G-V curves are fitted by Boltzmann function (solid lines) with V1/2 (24.1 ± 8.7 mV for WT+hβ4 and 33.6 ± 6.9 mV for N1159S+hβ4). The data are presented as mean ± SD (n=6).

### Identification of a benign variant p.E656A in the BK channel in a patient with epilepsy and molecular characterization

Whole exome sequencing analysis identified a novel A to C transition at codon 656 in a 30 year old USA patient with sever epilepsy (absence, atonic, myoclonic, tonic and other types), which results in the substitution of the glutamic acid residue for an alanine residue (p.E656A). The p.E656A is a variant of unknown significance which occurs at an amino acid residue that is highly conserved during evolution (Figure 8A). The patient is adopted and we could not test the parents to see if the variant is *de novo*. We recorded the voltage-current relationship of the WT BK channel and the mutant channel with the p.E656A variant (Figure 8C-F). No differences were observed between the mutant and WT channels with regard to the activation and current density of the BK channels in the presence or absence of the β4 subunit (Figure 8C-F). These data suggest that the p.E656A variant is a benign variant that is unlikely to cause epilepsy in the family.

**Figure 8.**
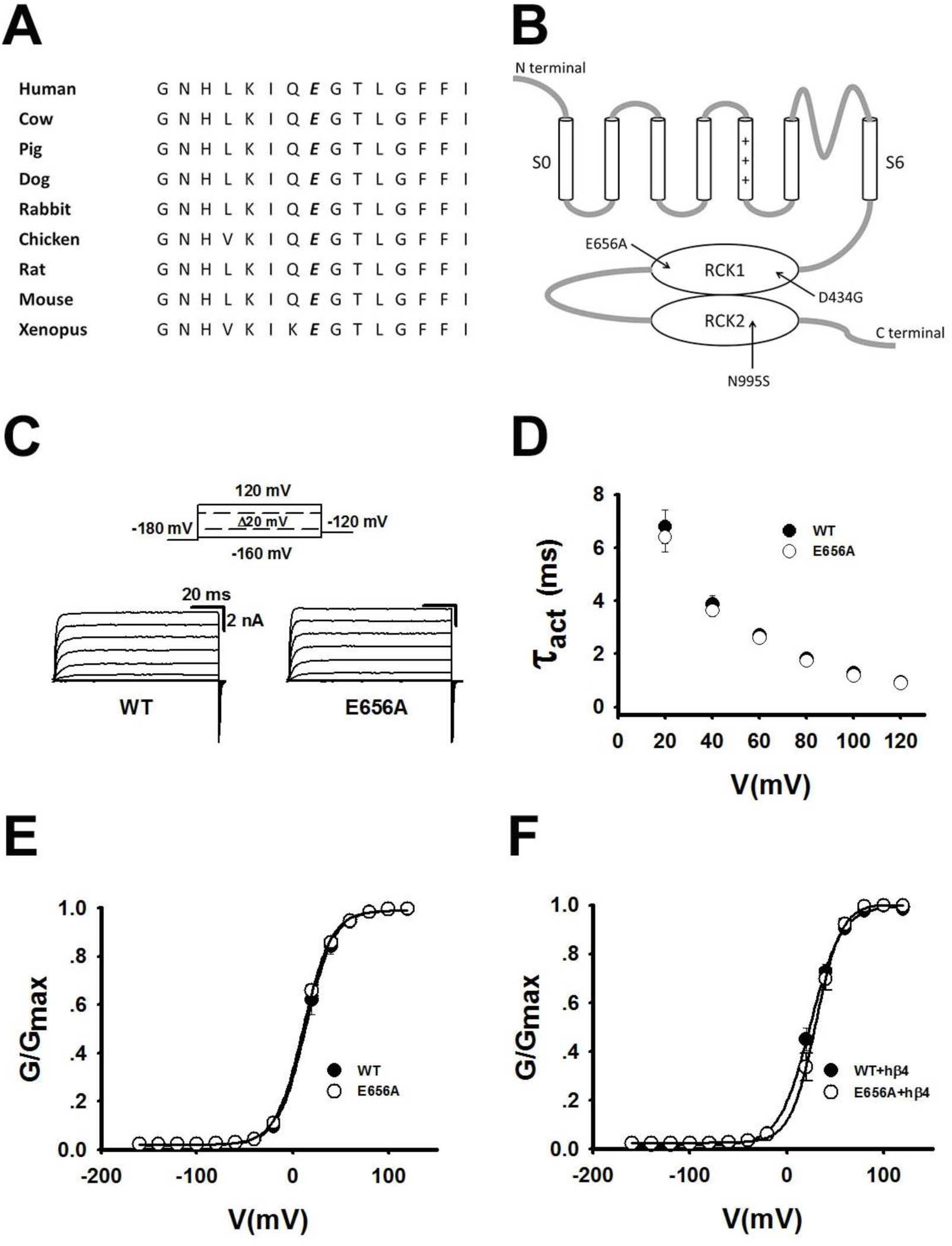
Whole exome sequencing identified a genomic variant E656A in the BK channel in a patient with epilepsy. (A) The E656 residue, shown in bold and italic, of KCNMA1 is evolutionally conserved among different species. (B) Structure of the BK channel with the E656A mutation indicated. D434G and N995S mutations are also shown. (C) Representative macroscopic currents of WT and E656A mutant BK channels from inside-out patches in the presence of 10 μΜ Ca^2+^ by the protocol as indicated. (D) Activation time constants of WT and E656A mutant BK channels at 10 μΜ Ca^2+^ are plotted against membrane potentials. (E) G-V curves of WT and E656A mutant BK channels at 10 μΜ Ca^2+^. All G-V curves are fitted by Boltzmann function (solid lines) with V1/2 (14.1 ± 6.6 mV for WT and 11.3 ± 3.1 mV for E656A). (F) G-V curves of WT+hβ4 and E656A+hβ4 at 10 μΜ Ca^2+^. All G-V curves are fitted by Boltzmann function (solid lines) with V1/2 (24.1 ± 8.7 mV for WT+hβ4 and 29.8 ± 6.3 mV for E656A+hβ4). The data are presented as mean ± SD (n=6).

## DISCUSSION

Our data in the present study demonstrate that mutations in the large conductance calcium-sensitive potassium BK channel can cause a common disease of epilepsy. This conclusion is strongly supported by both genetic and electrophysiological studies. The identification of a novel *de novo* mutation, p.N995S, in the BK channel in two independent patients with epilepsy demonstrates that p.N995S is high likely to be causative to epilepsy. Electrophysiological characterization of the p.N995S mutation demonstrated that it is a gain-of-function mutation that dramatically affects the functional properties of the BK channel (Figures. 2 and 3). Together, the genetic and functional electrophysiological data show that the p.N995S mutation in the BK channel causes epilepsy. Previously, we reported that the p.D434G in the BK channel causes a rare syndrome of co-existent generalized epilepsy and paroxysmal dyskinesia. The present study significantly expands the spectrum of clinical phenotypes caused by BK channel mutations from a rare syndrome to a much more common disease of epilepsy.

We have identified a novel molecular mechanism by which the p.N995S mutation causes epilepsy. The p.N995S mutation greatly increases macroscopic potassium currents from the BK channel by shortening the channel activation time and markedly shifting the G-V curve to the negative potentials by 59 mV at 10 μΜ Ca^2+^ (Figure 2). The p.N995S mutation can enhance the BK current by increasing the open probability and significantly prolonging the channel open dwell time probably by stabilizing the open state of BK channels at the single channel level (Figure 6). Increased potassium current by the mutant BK channel with the p.N995S mutation will result in the efflux of potassium, leading to a faster cell membrane repolarization or even hyperpolarization during the action potential. This shortens the duration of the action potential. In addition, the sodium channels that initiate the action potential are inactivated until the cell is repolarized. The recovery of sodium channels from inactivation is necessary for the generation of the next action potential. Faster repolarization or hyperpolarization resulting from the increased BK current by mutation p.N995S can enable a faster recovery of sodium channels so that the neurons can fire again at a higher frequency. Either the decreased action potential duration or an increased firing frequency of neurons will trigger the development of seizures and epilepsy.

Human genetics offers a novel strategy to identify drug targets for treatment of human diseases. Our demonstration that gain-of-function mutations of BK channels cause epilepsy suggests a strategy for the treatment of epilepsy. The link between the BK channel and epilepsy is also supported by the finding that the BK channel is most abundantly expressed in the cortex, hippocampus, piriform cortex, and other limbic structures, which are the brain areas involved in epilepsy (Knaus et al., 1996; Wanner et al., 1999). Because the increased BK current is a cause of epilepsy, BK channel inhibitors or blockers may be a novel strategy for the treatment of epilepsy. This is strongly supported by animal studies by Sheehan et al (Sheehan et al., 2009). In rats with injection of the gamma-aminobutyric acid (GABA)(A) antagonists picrotoxin or pentylenetetrazole developed seizures, but intraperitoneal injection of a BK channel blocker, paxilline, eliminated the tonic-clonic seizures, and reduced seizure duration and intensity in these animals (Sheehan et al., 2009). We found that treatment with 200 nM paxilline dramatically reduced the potassium currents from both WT BK channels and p.N995S mutant BK channels (Figure 2D and 2E). Some residual potassium current is still generated by the p.N995S mutant BK channels at the specific paxilline concentration (Figure 2D and 2E), which may be beneficial because a complete abolition of the BK current is too toxic for treatment. It is possible to develop a dose of paxilline that may block the BK current form mutant channels to a physiological level, which may be used to treat epilepsy. Future effort can also target the BK channel to develop specific inhibitors or blockers of the mutant BK channel activity suitable for the treatment of human patients with epilepsy.

It is interesting to note that the p.D434G mutation in the BK channel α-subunit causes a rare syndrome of co-existent generalized epilepsy and paroxysmal dyskinesia, whereas the p.N995S mutation causes epilepsy alone. There are two potential reasons for this observation. First, the penetrance rate of epilepsy and paroxysmal dyskinesia varies. Previously, we showed that in the family with the p.D434G mutation, 5/16 family members developed both epilepsy and paroxysmal dyskinesia (31%), 7/16 family members developed paroxysmal dyskinesia (44%), and only 4/16 family members developed epilepsy (25%) (Du et al., 2005). Therefore, the p.D434G mutation has a higher rate for predisposition to paroxysmal dyskinesia (44%) than to epilepsy (25%), whereas the p.N995S mutation has a higher predisposition rate to epilepsy (100%). Second, the functional effects of the p.D434G and p.N995S mutations are distinctly different. The p.434G mutation responds to both the voltage activation pathway and the calcium-dependent activation pathway. However, the p.N995S mutation responds to the voltage-dependent activation pathway alone and does not affect the calcium sensitivity. There is a possibility that the increased calcium sensitivity of the BK channel is associated with paroxysmal dyskinesia, but not with epilepsy, however, this speculation has to be vigorously tested by future experiments.

Our study identified a new functional domain responsible for voltage-dependent activation of the BK channel. Although the p.N995S mutation is located in the second RCK domain responsible for the calcium response of the BK channel, it did not affect the calcium-dependent activation of the channels because it still markedly enhanced channel activation under low calcium concentration and even when the calcium activation pathway is blocked by mutations (Figures. 4 and 5). On the other hand, the p.N995S mutation dramatically affected the voltage-dependent activation of the BK channel (Figures. 2 and 3). The p.N995S mutation enhanced the activation of BK channels by increasing the open probability and significantly prolonging the channel open dwell time (Figure 6), which may be due to a more destabilized close state so that the channel can open more frequently or be at a more stabilized open state of the channels so that the channels can open for a longer time. Based on the crystal structure of the cytoplasmic domain of the BK channel (Figure 5D) (Wu et al., 2010; Yuan et al., 2010), the N995 residue is located within the “turn” in the helix-turn-helix connector for the flexible interface that connects the RCK1 and RCK2 domains. Apparently, amino acid residues in the helix are important for the calcium-dependent activation of the BK channel. The N995 residue is in the turn of the helix-turn-helix connector, thus it is not involved in the calcium-dependent activation of the BK channel. On the other hand, the turn of the helix-turn-helix connector may be important for the overall molecular conformation of the BK channel through strengthening the interaction between the two RCK domains and other parts of the channel. The p.N995S mutation may alter the overall molecular conformation of the BK channel, weakening the close state of the channel or stabilizing the open state and making it open more frequently and for a longer period of time (Figure 6).

We found that the p.N995S mutation of the BK channel shifted the G-V curves to the negative potentials by more than 40 mV, but did not affect the calcium-dependent activation of the channel. The underlying molecular mechanism is unknown. In recent years, a series of leucine-rich repeat (LRR)-containing membrane proteins have been identified as BK channel auxiliary subunits, which were designated as the γ family of the BK channel auxiliary proteins (Yan and Aldrich, 2012; Yan and Aldrich, 2010). The γ subunits showed the tissue-specific expression patterns (Yan and Aldrich, 2012). The most notable, functional finding of the γ subunits is that they can significantly shift the G-V curve of BK channels to the negative potentials in the absence of intracellular calcium and did not affect the calcium sensitivity of BK channels (Yan and Aldrich, 2012). The effect of the γ subunits is remarkably similar to the effect of the p.N995S mutation in BK channels (Figures. 2 and 3). These data are consistent with the notion that the p.N995S mutation may affect the overall structure of the BK channel.

In the era of precision medicine, the data in our study clearly demonstrate the critical importance of functional characterization of genomic variants of unknown significance identified by whole genome sequencing or whole exome sequencing. In this study, we identified three genomic variants of unknown significance in the BK channel in patients with epilepsy. Both variants, p.N995S and p.N1159S, changed an asparagine residue to a serine residue. All three variants occur at the highly conserved residues in many species during the evolution. However, the p.N995S variant is a functional mutation that causes epilepsy, but the p.E656A and p.N1159S variants are not likely to cause epilepsy because of their lack of effects on BK channel function. Therefore, electrophysiological characterization of BK channel variants is required for distinguishing whether they have the potential to cause predisposition to a disease.

In summary, our data demonstrate that mutations in the BK channel α-subunit cause epilepsy. Our study identifies the first *de novo* mutation in the BK channel α-subunit associated with epilepsy. These results expands the BK channelopathy from a rare GEPD to a much more common disease of epilepsy. We have shown that the molecular mechanism for BK channel associated epilepsy involves the voltage-dependent activation, but not the calcium-dependent activation of the channels. We further show that electrophysiological characterization is required to determine a BK channel variant is causative to disease. We suggest that the BK channel is a novel potential target for the development of treatments for epilepsy.

## Materials and Methods

### Plasmid and mutagenesis

The human *KCNMA1* cDNA (GenBank accession number U23767) and *KCNMB4* cDNA (GenBank accession number AF207992) were cloned into a mammalian expression vector pcDNA3.1, resulting in expression constructs *KCNMA1-pcDNA3.1* and *KCNMB4-pcDNA3.1* for the α- and β4-subunits of the BK channel, respectively (kind gifts from Prof. Ding Jiuping at Huazhong University of Science and Technology).

We introduced mutations into the *KCNMA1-pcDNA3.1* expression construct using PCR based site-directed mutagenesis as described by us previously (Du et al., 2005; Huang et al., 2016). All wild type (WT) or mutant expression constructs were verified by direct DNA sequence analysis as described by us previously (Du et al., 2005; Huang et al., 2016).

### Cells culture and transfection

HEK293 cells were cultured in 24-well plates with DMEM supplemented with 10% fetal bovine serum (FBS) at 37°C in an incubator with 5% CO2. The cells were co-transfected with a *KCNMA1* expression construct and the pEGFP-N1 plasmid using Lipofectamine 2000 (Invitrogen), which generates a green signal to mark a transfected cell. During the transfection, the cells were incubated in OMEM, instead of DMEM supplemented with FBS. Five hours after the transfection, the OMEM was replaced with DMED supplemented with FBS. The transfected cells with a green EGFP signal were then selected for patch clamp recordings.

### Patch clamp recording

Patch clamp recordings were carried out 6-24 hours after the transfection as described by us and others (Cui and Aldrich, 2000; Du et al., 2005; Hou et al., 2013; Huang et al., 2016). All recordings were carried out with excised patches in an inside-out recording configuration. After obtaining a patch, we moved the electrode tip to a blank area of the chamber and then let the electrode tip touch the bottom of the chamber to expose the inside face of the patch to the bath solution so that the inside-out mode was established. Patch pipettes were pulled from borosilicate glass capillaries using Micropipette Puller P-97 (Sutter Instrument) with resistance of 2-5 megohm when filled with the pipette solution. Experiments were carried out using a Multiclamp 700B amplifier with a Digidata 1440A A/D converter and pClamp software (Axon Instruments). Recordings of currents were typically digitized at 20 kHz and low-pass filtered at 2 kHz.

For single channel recordings, the patches that contain only one channel were selected, and the recordings were digitized at 50 kHz and low-pass filtered at 5 kHz as described by us and others (Cui and Aldrich, 2000; Du et al., 2005; Hou et al., 2013; Wu et al., 2008). During the recording, different concentrations of the calcium solution was applied onto the membrane patches via a perfusion pipette containing eight solution channels. All experiments were performed at room temperature (22-25°C). The solutions used for patch-clamping include:

> Pipette solution containing (in mM): 160 MeSO_3_K, 2 MgCl_2_, 10 HEPES. Nominal 0
>
> μΜ Ca^2+^ solution containing (in mM): 160 MeSO_3_K, 5 EGTA, 10 HEPES.
>
> 1 μΜ Ca^2+^ solution containing (in mM): 160 MeSO_3_K, 5 EGTA, 3.25 CaCl_2_, 10 HEPES.
>
> 10 μΜ Ca^2+^ solution containing (in mM): 160 MeSO_3_K, 5 HEDTA, 2.988 CaCl_2_, 10
>
> HEPES.
>
> The pH of all the solutions was adjusted to 7.0.

## Data analysis

Recorded data were analyzed with Clampfit (Axon Instruments) and SigmaPlot (SPSS) software programs as described by us and others (Cui and Aldrich, 2000; Du et al., 2005; Hou et al., 2013; Wu et al., 2008). For the analysis of macroscopic currents, we utilized the G-V relationship to analyze the kinetic property of the BK channels. G-V curves for activation were fitted by the single Boltzmann function with the formula: G/G_max_ = 1/(1 + exp((V_1/2_ - V)/κ)) where V_1/2_ is the voltage where the conductance (G) is half of the maximum conductance (G_max_) of the channels and κ is a factor indicating the steepness of the G-V curve. We fitted the current traces by single exponential function to obtain the activation time constant.

For the analysis of single channel currents, we analyzed the open probability (P_o_), single-channel conductance and open dwell time of the BK channels as described by us and others (Cui and Aldrich, 2000; Du et al., 2005; Hou et al., 2013; Wu et al., 2008). We detected the events in the recordings with half-amplitude threshold analysis. The P_o_-V curves were also fitted by single Boltzmann function. We fitted the amplitude histograms by Gaussian function to acquire the current amplitude of the single channel, and then the quotient of the current amplitude and given voltage was the single-channel conductance. We analyzed the open dwell time of the channels by fitting the open dwell time histograms by exponential function.

## Human subjects

The study subjects were patients with epilepsy and their family members. This study was approved by the Institutional Review Boards (IRB) on Human Subject Research at the Cleveland Clinic and the Ethics Committee on Human Subjects at Huazhong University of Science and Technology and other local IRBs. Written informed consent was obtained from the study subjects according to the IRB policies.

Human genomic DNA samples were isolated from peripheral blood leucocytes using standard protocol as described by us previously (Wang et al., 2016; Wang et al., 2010; Xu et al., 2014).

Whole exome sequence and targeted sequencing were performed as described (Wang et al., 2016; Wang et al., 2010; Xu et al., 2014).

## Statistical analysis

All data were shown as mean ± SD. A student’s t-test was used for comparing the means from two groups of variables. A *P* value of 0.05 or less was considered to be statistically significant.

## Acknowledgements

We are grateful to Dr. Johannes Lemke for help and assistance and Dr. Jiuping Ding for the gift of the expression construct for *KCNMA1* and *KCNMB4*. We thank other members of Wang laboratory for help and assistance. We greatly appreciate all study subjects for their participation in this project. This study was supported by the China National Natural Science Foundation Key Program (31430047 and 81630002), Chinese National Basic Research Programs (973 Programs 2013CB531101), Hubei Province’s Outstanding Medical Academic Leader Program, Hubei Province Natural Science Key Program (2014CFA074), the China National Natural Science Foundation grants (91439129, NSFC-J1103514, 81370206), NIH/NHLBI grants R01 HL121358 and R01 HL126729, a Key Project in the National Science & Technology Pillar Program during 395 the Twelfth Five-year Plan Period (2011BAI11B19), Specialized Research Fund for the Doctoral Program of Higher Education from the Ministry of Education, and the “Innovative Development of New Drugs” Key Scientific Project (2011ZX09307-001-09).

## Author Contributions

X.L. and Q.K.W. designed the study. X.L., S.P., N.J.O., F.M.S., E.J.K., M.W., Q.C. and Q.K.W. performed experiments, acquired, analyzed and interpreted data. X.L. and Q.K.W. drafted manuscript. M.W., Q.C. and Q.K.W. edited and revised manuscript. M.W. S.P., F.M.S. and Q.K.W. supervised the clinical and basic aspects of this project.

## Ethics

This study was approved by the Institutional Review Boards (IRB) on Human Subject Research at the Cleveland Clinic and the Ethics Committee on Human Subjects at Huazhong University of Science and Technology and other local IRBs. Written informed consent was obtained from the study subjects according to the IRB policies.

